# Current systems biology approaches in hazard assessment of nanoparticles

**DOI:** 10.1101/028811

**Authors:** Friederike Ehrhart, Chris T. Evelo, Egon Willighagen

## Abstract

The amount of nanoparticles (NPs) in human environment is increasing. The main sources are the increased introduction in consumer products and air pollution (diesel exhaust). It is meanwhile common knowledge that NPs behave differently as bulk material because of their nano-size. This leads in general to a higher reactivity and some other changed properties, *e.g.* solubility, surface potential, conductivity, and, to different effects on biological systems. The main impacts of NPs on a cellular and organism level are meanwhile well known: release of toxic ions, increased oxidative stress, and inflammation. Beside these, there is increasing evidence that NPs, especially in low dose/long exposure scenarios, affect biological systems in a broader way, interact with drugs, and exacerbate the effects of diseases. To investigate these effects systems biology approaches are the method of choice. This review summarizes the state of the art of nanoparticle effects on cells and organisms and demonstrate the add value of systems biology investigations to NP hazard assessment.

Abbreviations: nanoparticle (NP), carbon nanotube (CNT), single walled carbon nanotube (SWCNT), multi walled carbon nanotube (MWCNT), ADME (adsorption, distribution, metabolization, excretion), PAMAM (poly(amidoamine)),

## 1. Introduction

Nanoparticles (NPs) are particles defined by their size. The most commonly used definition for NPs is being between 1 and 100 nm in at least one dimension (definition according to ISO^1^). That size gives them special properties which are often quite different from bulk material and makes them interesting for research and application. In general, NPs have a drastically increased surface/volume ratio which increases their chemical reactivity. The most abundant NP in the world is assumed to be carbon black from diesel motor combustion exhaust, while the most abundant engineered NP in industry, consumers products respectively, are SiO_2_, TiO_2_ and Ag NPs. Medical applications focus on magnetic NPs or NPs used as drug delivery systems. In the past decade it has become common knowledge that nanomaterials are special and must not be treated like bulk material. Risk assessment is basically collecting data (and producing if necessary), calculating and understanding the risk of a defined entity or action. The first step to assess the risk of any chemical substances is to gain prior knowledge about dose and exposure dependent effects: the hazard assessment. What makes the assessing of NPs' hazard so difficult is the strongly size and surface dependent behavior.

Particles of the same material source but small differences in size, surface (coating), charge or solvent components may exhibit different properties^2^. Systems biology is a holistic approach to model complex biological systems using mathematical and computer science methods. Basis of systems biology are the single components and entities of a biological system: genes, transcripts, proteins, and metabolites (see Figure 1). A good understanding and a description of the single entities and how they are interacting within a biological system is crucial for systems biology approaches. Based on this, time or incident dependent changes, *e.g.* exposure of NPs, in the system can be investigated. The interactions and relations between the entities are commonly described in biological pathways and networks. The state-of-art measuring technologies like genome sequencing and high throughput measurements enables the researchers to study the overall behavior and phenotype of a biological system in a defined setting. So, besides “listing the parts of an airplane” the scientific community proceeds to study the system as a whole^3^. This thorough understanding finally allows prediction of behavior using computational models.

**Figure 1.**
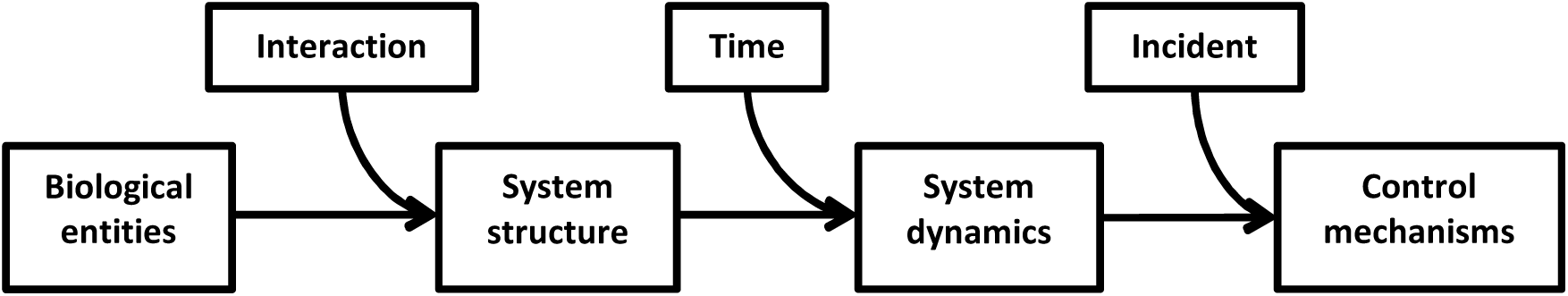
Scheme of systems biology. Biologic entities like genes, proteins or metabolites are connected by interactions forming the structure of the system. This structure alters with time and reacts to incidents demonstrating systems dynamics and control mechanisms.

In this review we focus on the actual status of hazard assessment of NPs and the added value of using systems biology tools and methods in the assessment. We excluded NPs for therapeutical use because cost-benefit analysis for drugs is differently evaluated.

## 2. NP toxicology

NP hazard assessment is in a way different from “ordinary” chemicals as NPs derived from the same bulk material may react differently according to their size, shape, surface, or other properties. The precise characterization of a NP before starting hazard assessment experiments requires therefore a detailed dossier including the following information:

- **Physico-chemical properties**: particle morphology (size, shape and aspect ratio), water solubility, surface area, chemical composition (including impurities and crystallinity), surface chemistry, surface charge and surface hydrophobicity, impurities and ions, storage, dissolution and agglomeration information;
- **General biologic activity**: dissolution rate (in a given scenario), surface reactivity, abiotic ROS generation, redox and photocatalytic activities, dispersibility, corona formation^4,5^, surface area *in situ,* and biopersistance;
- **Consistent NP production** batches indicating a stable, ideally a validated, production process.

### a. Uptake and distribution

**How much NPs are we exposed to?** In Europe the daily limit for the so called PM10 (particles with less than 10 μm) is 50 μg/m^3^ and may not be exceeded more than 35 times per year. For the smaller fraction of PM2.5 (less than 2.5 μm) there is a limit of 25 μg/m3. Except for air pollutants, it is quite difficult to measure the actual amount of engineered NPs in *e.g.* soil and surface water so exposure modeling is currently the best guess. Taking the yearly production and lifecycle of engineered NPs as a basis, Gottschalk and coworkers^6^ modeled the environmental concentration in surface water, soil, sewage treatment effluents etc. and predicted concentrations ranging from 0.003 ng/l (fullerenes) to 21 ng/l (TiO_2_ NPs) for surface waters and from 4 ng/l (fullerenes) to 4 μg/l (TiO_2_ NPs) for sewage treatment effluents. The yearly increase of NPs on sludge-treated soil was estimated between 1 ng/kg (fullerenes) and 89 μg/kg (TiO_2_ NP). Another risk assessment study from Maynard and coworkers^7^ calculated an amount of 0.039 mg SWCNT (Single Walled Carbon Nano Tubes) deposited in the human lung during an eight hour working day assuming 53 μg/m^3^ particles in the air. Uptake of NP via food (mainly TiO_2_ and SiO_2_) is estimated to be more than 10^12^ particles/day^8^ (for more information about natural and anthropogenic NP sources and exposure see Buzea et al.^9^ and literature cited there).

**NPs enter the body** via dermal, nasal, ocular, respiratory and gastrointestinal route^10,11^. Experiments on dermal uptake showed that silver NPs are able to penetrate porcine skin to 10–18 μm in 16 h^12^. Reviewing the effect of sun screen particles (mainly ZnO and TiO_2_) the authors propose the following pathways for NPs entering through (healthy) skin: The transappendagael (sweat pores, hair follicles, skin glands), paracellular or transcellular route^13^. While some authors of exposure studies did not find TiO_2_ others found it in all skin layers and also in secondary organs such as lung, brain and spleen (see ^13,14^ and literature cited there). However, the uptake by skin or contact with eye fluid is very small and most studies focus on the bulk uptake via gastrointestinal and respiratory system. NPs are difficult to track in an organism. Some uptake detection limits were given in van der Zande et al.^15^: for silica using HDC-ICP-MS: 300 mg/kg tissue, SEM-EDX: about 100 mg/kg tissue. Tracking experiments showed *e.g.* that airborne NPs were translocated within 4 – 24 h from the primary uptake site – the lung epithelium - to interstitial sites in the respiratory tract and from there to all other organs. A partition of NPs is transported via bronchial mucus from bronchia epithelium into the gastrointestinal tract^16^. Experiments demonstrated that inhaled NPs can be taken up directly from nasal skin into nerve cells of the olfactory bulb and were found again in brain cells ^17,18^. After injection of TiO_2_ NPs in the blood circulation system accumulation was found in liver, lungs and spleen, possible due to resident phagocytotic cells there^19^. A feeding study reported a significant uptake and accumulation after 28 days of exposure in liver, kidney and spleen which did not change after 84 days except for the spleen which still increased the amount of silica NP^15^. The uptake rate and bioavailability of polystyrene NPs ranged between 0.2 and 1.7 % *in vivo,* which was much less than the values found for *in vitro* uptake: 1.6 - 12.3 %^20^.

**ADME** (adsorption, distribution, metabolisation and excretion): Concerning pharmacokinetics, some authors speak about “clearance” of NPs and mean the uptake of NPs by phagocytic cells, namely the clearance of NPs from blood stream not the clearance from the organism which shall be dealt with here^21^. The biopersistance of NPs, their lifetime in an organism, depend strongly on their material. Protein, biodegradable polymers, DNA and soluble metal particles are generally dissolved, metabolized or cleared from the organism within a material specific time^22^. Talking about non-degradable, biopersistant NPs, very small NPs (below 10 nm) are cleared quicker via the kidneys and have shorter blood half-lives than larger particles (20 – 100 nm) because of the size exclusion filter of the glomeruli, which is about 5–7 nm or 67 kDa. Dufort et al.^23^ stated that particles with less than 40 kDa will be cleared up soon while large NPs (up to 500 nm) will be retained in the interstitial space (of tumors) due to lack of drainage. Small amounts of NPs in tissue are difficult to prove, nevertheless, it is assumed that they can reside over long periods in the body, *e.g.* over two years^22^.

**Uptake in the cell:** NPs have the same dimension as large biological macromolecules like proteins, RNA or vesicles. Uptake has been observed in all cell types and currently there is no publication available where the rejection of a truly nano-sized NP by a cell (*in vitro!)* has been reported. The uptake rate, nevertheless, may differ greatly and depends on different factors like size and surface properties^10^. Phagocytosis specialized cells like macrophages take up Au NPs using the macrophage scavenger receptor A^24^, while mannose-coated NPs are taken up via the mannose receptor-mediated phagocytic pathway^25,26^. Another study indicated the complement receptor-mediated internalization pathway to take up lipid nanocapsules^27^ and fullerenes were taken up by Fc⍰R^28,29^. Surface coating with hydrophobic polymers like PEG were used to reduce NPs uptake in phagocytosis specialists and to allow them (the NPs) to spread more in the test organism and be taken up by non-specialized cells^24^. NP binding rate on plasma is an indicator how fast they will be cleared by phagocytotic cells^30^. Beside phagocytosis, pinocytosis/macropinocytosis, clathrin-mediated endocytosis, caveolin-mediated endocytosis, and non-clathrin-mediated endocytosis are all possible entrance pathways for NPs^31,10,32^. *Vice versa,* three different exit routes are described: Non-vesicle related secretion, lysosome secretion and vesicle related secretion^33,34^. Very small NPs may enter the nucleus through the pores: gold NP of 5 nm, *e.g.,* were found in the nucleus of human fibroblasts^35^. Simulation studies by Lai et al. 2013^36^ showed that CNTs can pass biological membranes due to their hydrophobic nature and affect the membranes stability.

**Intracellular fate:** Most NPs will end up at some point in lysosomes. There, they are confronted with a quite hostile environment: acidic pH, hydrolytic enzymes, H_2_O_2_ etc. Non-biopersistant NPs are dissolved in this stadium and released in the form of molecules or ions into the cell metabolism. Biopersistant ones will stay in the lysosome until fusion with autolysosomes or other vesicles, connection to ER and Golgi, or dissolution of the vesicle and enter the cytoplasm. Using scanning transmission X-ray microscopy Ahlberg et al. found silver NPs to be accumulated in the perinuclear region of human mesenchymal stem cells and they found some evidence for NPs being aggregated in lysosomes^37^. NPs are able to cross biological barriers like the placenta^38,39^ and blood-brain-barrier (e.g. via nasal uptake), which makes them valuable tools for drug delivery but it means also that neither fetuses nor brain nor any other tissues are protected^18^.

### b. NP effects *in vitro*

In general, getting valid hazard assessment data about NPs from *in vitro* studies requires awareness of the following pitfalls (summarized in ^40^ and ^41^): Due to corona formation and specific solubility in physiological solutions (*e.g*. presence of organic acids which withdraws metallic ions) the NPs need to be carefully characterized in each experimental setting. This includes the careful observation of the protein environment which contributes to the corona. The choice of cell type also strongly influences the outcome of the experiment. In general, the following effects were observed after exposure of cells to nanomaterials:

- Cytotoxic effects of toxic ions released by NP;
- Increased production of radical oxygen species (ROS) causing oxidative stress:

- Induction of necrosis, necroptosis, apoptosis, increased autophagy;
- DNA damage, genotoxicity, increased mutation rate;
- Membrane damage;
- Cytoskeleton and cell architecture defects

- Lysosome overload which results in lysosome damage;
- Frustrated phagocytosis.

Metal or metal oxide NPs generally **release ions**. It is therefore not surprising, that NPs made of toxic metals like cadmium^42,43^ or lead^44,45^ are toxic to cells. Heavy metals are used for quantum dot NPs but traces of these or other ions may be found as catalyst residues, production by-products, and reaction side products in other NP suspensions, too.

Silver inhibits bacterial growth and silver NP are used to deliver ions, *e.g.* to wound dressings, socks or refrigerator surfaces. The antimicrobial mechanism of silver is explained by inhibition of DNA replication and protein inactivation and there are *in vitro* studies revealing an toxic effect to mammalian cells via oxidative stress^46^ or apoptosis induction^47^, too. It remains to be elucidated whether Ag NPs are more toxic than an equivalent amount of Ag ions because of contradictory study effects (see Munger et al.^48^; for a review of toxic silver effects in general see also ^49^). Astrocytes are quite resistant to toxic metal ion effects caused by silver NPs possibly due to their metal homeostasis equipment^50^. Effects like this may explain why some cell types (or some species) are more vulnerable to ion toxicity than others just like reported in Brunner et al. who found different cytotoxicity rankings of ion-releasing NPs for different cell types^51^.

The active **production and release of reactive oxygen species** (ROS) (*e.g*. superoxide radicals (O_2_•-), hydroxyl free radicals (•OH)), or the reduced detoxification (decrease of superoxide dismutase or catalase), the depletion of antioxidants, respectively, is a general cellular response to stress factors and can be activated by a broad variety of factors. Cells are usually able to get rid of ROS which are produced during normal metabolic function. Increased production because of stress factors overloads the disposal system and ROS react freely, oxidizing proteins, lipids, and DNA. Changes in oxidative stress response were regularly observed in cells *in vitro* after exposure of NPs which led to *in vivo* local and systemic inflammation response, DNA damage and tissue injury^39 52–54^. Increased oxidative stress is known to trigger a variety of cellular pathways, interfering with cell cycle and finally leading to cell death^32^. Apoptosis induction, oxidative stress and mitochondrial damage were reported for CNT^55^, ZnO^54^ and Au NPs^56^. Antioxidative stress pretreatment reduced in some cases the damage of CNT^57^ or TiO_2_^58^ NPs.

The **cytoskeleton** plays an important role in regulation and trafficking of cellular vesicles like lysosomes or autophagosomes^59^ and beside cytoskeleton this function was also impaired in MWCNT (multi walled carbon nano tubes) treated keratinocytes^60^. Fullerenol NPs highly impaired the cytoskeleton, causing actin filament disruption and clumping, and induced autophagic vacuole accumulation while there was in that case no oxidative stress observed^61^. SWCNT do the same damage to the cytoskeleton of fibroblasts^53^ and endothelial cells^62^. Lysosomes suffer from NP overload which impairs their function especially concerning autophagy^31^. CNT decreased cell adhesion by reducing expression of cell adhesion related proteins leaving the cells less attached to the surface and adopting a spherical shape due to missing focal contacts^53^. A more specific reaction towards fibrous NPs is the so called “frustrated phagocytosis”. This effect was first observed when lung macrophages were confronted with asbestos and similar effects were predicted and proven for fiber shaped NPs, too^63–65^. While short fibers are successfully cleared and found in the cytoplasm, long fibers resist to macrophage clearance causing cellular stress^63^.

2010 Hussain and co-workers^66^ reported **apoptosis** of cells after NP exposure and they proved that there are two different pathways for apoptosis activated by two different kinds of NPs. One year later Ma et al. ^67^ reported investigations of water-soluble germanium NPs, which caused **necrotic cell death**. They found that the damage can be attenuated by blocking the transduction of necrotic signaling pathway which proofed that necrosis not apoptosis was responsible for cell death in their experimental system. A recent study from Andon and Fadeel^68^ summarized four ways of more or less regulated cell deaths induced by NPs: apoptosis, autophagic cell death, regulated necrosis (necroptosis) and necrosis. NPs are in several studies found to activate different lethal pathways in a size, dose and type dependent way^56,69,70,71^.

**Epigenetic changes** were, as reviewed in Stoccoro et al.^72^, reported for several NPs: airborne NPs from diesel exhaust changed methylation and acetylation patterns in various biological models, and miRNA expression was affected, too. Engineered NPs like SiO_2_, Au, or MWCNTs were found to cause *i.a.* global DNA hypomethylation, chromatin condensation and deregulation of miRNA expression.

### Nanomaterial effects *in vivo*

*In vitro* studies generally confront cells with much higher NP loads than cells ever meet *in vivo*^20^. Whereas similar effects were measured – *e.g.* oxidative stress – the levels were lower *in vivo* than *in vitro^73^* which is due to the low uptake rate of NPs (see 2.a). For a summary of correlation studies between *in vitro* and *in vivo* effects see Maurer-Jones and Haynes 2013^74^.

The **respiratory tract** is the dominant entrance site, the first site of action and reaction and therefore the most abundant investigation model for airborne NP effects. A lot of animal experiments showed that the first reaction of lung epithelium to a single dose of NPs was an acute but reversible inflammation whereas the severity depends on the amount and type of NP. If chronic responses were monitored, e.g. after single or repeated dose, authors reported fibrosis or lung tumors^75^. The acute inflammation showed typically symptoms like fever, dyspnea, attraction of immune cells, vascular and epithelial permeability, hypercoagulation, hypofibrinolysis and immune modulation^75^. Metal fume fever^54,76^ is associated with air pollution in industrial production sites where particles containing Zn, Fe, V and Ni and their oxides are released. Cerium pneumoconiosis, or rare earth metal pneumoconiosis, was observed with peoples exposed to metal gases of that kind which also contains NPs^77^. Lung macrophages are stimulated by NPs and react with inflammatory response in general^78^. Long term macrophage stimulation, e.g. suffering from frustrated phagocytosis caused by fibrous NPs, and inflammation produce a carcinogeneous environment leading to fibrosis and cancer^79^.

Phagocytotic cells of **the immune system** are one of the first ones to pick NPs up, once they crossed the epithelial layer. Recognition of NPs by immune system is facilitated by the surface properties of NPs, which attract protein to form a corona^5^. A strong reaction of lung macrophages towards particulate matter was observed to lead to granuloma formation by production of an inflammatory environment^75^. Some NPs like PLGA^80^ (poly(lactic-co/glycolic acid)) are able to stimulate the complement system which may lead to severe side effects if used as drug delivery systems^81^. There is also a current discussion of a possible connection between high NP uptake and Crohn's disease or ulcerative colitis (both chronic gut inflammation)^8^ and some autoimmune diseases^9^. Most studies confirm a pro-inflammatory and immune system stimulating effect of NPs but also **immune system suppression** was observed, *e.g.* after MWCNT inhalation a dose dependent depression of T-cell dependent antibody response was observed in mice^82^. PAMAM-dendrimer NPs were discovered to have an anti-inflammatory effect if the shell contained large amine- and hydroxyl-terminated dendrimeres. Some other NPs act as an anti-inflammatory agent by intrinsic properties: Cadmium is generally known to suppress the immune system by reducing the subpopulation of CD4+ and CD8+ cells in a dose dependent way^42^. A summary of NP effects to the immune system is given in Ilinskaya et al.^83^ and Zolnik et al.^84^.

Epidemiological data revealed that high air pollution in general triggers **myocardial infarcts** in a population and this is mostly referred to the particular matter containing NPs^85^. Coronary heart disease, the development of cardiovascular plaques, and infarcts are triggered via inflammation and metabolic/oxidative stress mediators (like c-reactive protein, CRP)^86,87^. Chronic exposure to NP activated mitochondrial aortic alterations which contributed to plague formation and coronary heart disease^88^. There is some evidence that increased Ce (which is used as a fuel additive and released after combustion in the form of CeO_2_ NPs) concentrations increase the risk of cardiac infarcts^89^. An exposure study on 62 patients which got orally 10 or 32 ppm silver NPs revealed no significant heart arrhythmia signals during NP uptake^48^. Another controlled short term exposure study with healthy and coronary heart disease patients did not show an increased incidence of heart arrhythmias^90^ during the experiment. But these findings may be due to the short exposure and observation time and the long term inflammatory process of disease development.

**The liver** is an accumulation site for gastro-intestinally or tracheally delivered NPs. Due to resident phagocytic cells NPs are taken up easily and they are after the lung the primary accumulation sites for NPs in an organism. *In vitro* studies provided some information about oxidative stress induced by NPs to hepatocyte cell lines like HepG2^46^ or BRL 3A^91^. An animal study showed that consecutive feeding of rats with TiO_2_ resulted in suppression of hepatic glutathione levels and triggering of inflammatory response by activation of macrophages. This observed liver toxicity of TiO2 was partly reversible by parallel feeding of antioxidants^58^. Repeated injections of TiO2 NP in the peritoneum of mice resulted in severe histopathological changes and liver cell apoptosis. The liver function was impaired and a general upregulation of inflammatory cytokines was observed^92^. Oxidative stress and apoptosis were, too, the cause of liver toxicity of ZnO NP fed to mice^93^. These effects seems to be strongly dose, exposure time and administration pathway dependent because the study of van der Zande et al. found no pathologic changes in liver tissue after feeding low amounts of nano-silica to rats after 28 days, but a slight increase of liver fibrosis after 84 days^15^.

NPs inhibit **stem cell** differentiation and other reproductive and developmental processes as reviewed by Ema et al. ^94^. Li et al. summarized the known **effects of NPs to sensitive populations**, namely the neonate, pregnant, diseased, and old, and resumed that they generally suffer under exacerbated symptoms while healthy individuals often are unaffected^38^.

### Add value of omics approaches

The main toxic effects of NPs to cells and organisms are yet quite well understood but there is a systemic level which cannot be addressed by these methods. Using the gain by high throughput data we are currently able to include complex information about biologic networks to get a more detailed picture of NP induced incidents. The main advantage of omics approaches is the impartiality in which the experimental read-out is performed. By selecting a read out the researcher sets the limits of the test, *e.g.* an assay on oxidative stress induction would not reveal additional information about mitochondrial metabolism or chromatin methylation. Tools like the Generic Gene Ontology Term Mapper^60^ *e.g.* help to discover long-range effects of MWCNT by revealing a broad variety of pathways and effects: apoptosis, cytoskeleton, membrane trafficking/exocytosis, metabolism, tox/detox, protein degradation, and stress proteins/chaperons. This load of information would have taken more time and animal lives to be discovered and investigated on the single experiment level. A broader investigation of pathways involved already led to the discovery of cross-toxicities as shown by Triboulet et al. ^95^ between Cu NPs and drugs, which interfere with the glutathione biosynthesis.

Omics approaches contribute to a better understanding of NP activated (or depressed) pathways and allow an improved prediction of NP effects. The amount of data from omics experiments usually covers most parts of cell metabolism and toxic phenotypical effects. So, *in vivo* responses can be extrapolated from *in vitro* experiments, long term effects from short term experiments – *e.g.* cell division in lung epithelium^96^, and low dose exposure experiments may give additional information about effects of high doses^97^. Classification of NP toxicology according to the amount and kind of pathways activated and extrapolation of those to new designed NPs contribute strongly to the Safe-by-Design approach of NP development^98^.

The strength of –omics approaches are in the field of complete picture description and their limits lie in the detail: the precise, quantitative description of single read outs. Using *e.g.* a transcriptomics approach, the result will not reveal whether an increased transcription finally led to an increased protein concentration and a change in metabolism at all. A possibility to cover that would be either to use different –omics approaches together, *e.g.* using proteomics experiment to verify a changed transcriptomics image, or to verify the interesting findings by chosen classical experiments.

## Systems biology tools and methods / Data analysis in biological context

### Data acquisition

Recent technical developments allow laboratory scientists not only to look at the expression of a single or a few entities of interest but up to a few hundred or thousands. Due to the improvement of high throughput analysis methods like DNA and RNA sequencing, microarrays, mass spectroscopy (MS) or chromatography (HPLC, MALDI-TOF) methods biologists can gain information about a few hundred up to ten thousands of data points per sample. This allows us to gain a more detailed picture of the metabolic processes involved in the experiment.

The first step of an analysis is to identify the entities which are changed in a certain experimental setting. To gain that information, the *in vitro* or *in vivo* samples must be treated according to the experimental setup and processed. In case of nanomaterial studies, the standard investigation procedure is to confront cells or animals with NPs and examine their reactions.

The technique of DNA microarrays *e.g.* has its origins in Southern blotting, a technique to detect short specific DNA sequences in a sample. This principle is miniaturized and expanded to a few thousands of DNA or RNA sequences on a microarray which are used to analyze single nucleotide polymorphism (SNP), gene expression profiles of cells or investigate epigenetic information (Chromatin immunoprecipitation on Chip). Similar miniaturized and improved detection systems are also available for metabolites and proteins.

The raw and processed experimental data are often requested to be deposited on a public available database like ArrayExpress^99^ from European Bioinformatics Institute (www.ebi.ac.uk/arrayexpress EBI, GB) or Gene Expression Omnibus (GEO) from the National Center for Biotechnology Information (http://www.ncbi.nlm.nih.gov/geo NCBI, USA)^100^. This creates a stock of experimental data for future re-investigation of existing data sets with new or improved methods, to compare new with old datasets for control and it is easy to check if an experiment was already done before. Multiple datasets – *e.g.* from different laboratories - proving and predicting the same results increase in general the relevance of the result.

### Data visualization and biological interpretation

In most cases, -omics analyses result in a list of up and down regulated genes. It would be much paperwork for the interested scientist to work through this list and find the relevant information to answer research questions like “Are inflammatory related processes affected in this experiment?” Therefore, data analysis should include association and visualization methods. Association tools help *e.g.* to find the connections of genes/metabolites with pathways or phenotypes. Visualization makes analysis and interpretation easier or even possible especially when it comes to highly interconnected data points.

Example for association studies are *e.g.* gene ontology or pathway analysis. These techniques help investigating association enrichment in genes or grouped genes (pathways). The Gene Ontology project provides consistent description of the biological functions. Gene Ontology analysis finds enriched biological functions for the list of changed genes in the experiment. Pathways, on the other hand, provide a detailed, visual representation of a biological process. Pathways are categorized into gene regulation, signaling, and metabolic pathways. Pathway analysis maps genes involved in a certain experiment to known pathways and determine which pathways are over- or underrepresented. By revealing the pathways affected during an experiment the results provide mechanistic insights into the underlying biology. Both Gene Ontology and pathway analysis reduce the complexity from thousands of genes to a smaller set of biological processes and functions allowing a quicker and more detailed analysis. A model systems biology workflow example is given in Kutmon et al. ^101^.

### Databases

The basis for every further manual or automated data analysis is information: efficiently organized, safely stored, and online available information. Data bases store information about biological entities like genes, proteins or metabolites and makes them available for manual and automated information request. The most famous examples are Ensembl (http://www.ensembl.org) and NCBI Gene (http://www.ncbi.nlm.nih.gov/gene) which provide information about genomes, genes and transcripts. UniProt (www.uniprot.org) contains information about proteins and ChEMBL (https://www.ebi.ac.uk/chembl) is a database for metabolites. Links between single entities are deposited in interaction databases like the protein-protein interaction database IntAct (www.ebi.ac.uk/intact) while WikiPathways (http://www.wikipathways.org), Reactome (http://www.reactome.org/) and KEGG (http://www.kegg.jp/) focus on biological pathways. Functional information like ontology databases, used for e.g. term enrichment analysis, is found on BioPortal (http://bioportal.bioontology.org). The information available for biological (and chemical) entities is quite well integrated in the scientific community (as *e.g.* shown by the broad variety of analysis and visualization tools which require access to those databases), but there is currently little information available about NPs and their special properties. Specialized databases for NPs and nanomaterial investigation are *e.g.* nanowerk (http://www.nanowerk.com) and nanomaterial registry (https://www.nanomaterialregistry.org). A computational infrastructure for toxicological data management for NPs, including ontologies, is currently developed in the eNanoMapper project^102^.

## Hazard assessment of NPs

NP risk assessment is most important for manufacturers, laboratory personal and consumers. An overall risk assessment, however, should include knowledge about:

- severity of hazard (or hazard assessment);
- probability of incident or exposure;
- probability of detection (of an incident/ exposure);
- possibility of medical treatment.

In this section we discuss the current status of nanotoxicological hazard assessment and propose a workflow how it could be improved for new materials.

### Conclusion on NP effects

Currently it seems that neither classical experiments nor–omics approaches find NP specific toxicity effects. For a specific particle there may be debates about whether its toxicity is related to impurities, release of ions, or nano size, or all of them. Consensus is that the nano size itself evokes unspecific stress reactions leading to oxidative stress with all its main and side effects. In addition, there are NP type, tissue or cell specific reactions after exposure. So, there is common agreement that NPs do cause damage in cells and tissues and must therefore not be treated like bulk material and there is experimental evidence for certain parameters to contribute more to toxic effects than others. The questions to be solved are: “to what extend can we predict toxicity of new NPs according to existing data?” “which experiments do we have to execute to assess NP hazard?” and finally, “which dose is acceptable for short term, long term, or maximum workplace exposure?”

### Current status of NP risk assessment

There are several attempts to classify nanomaterials *e.g.* by their main component (carbon or metal based materials), physico-chemical properties^103,104^, by their size or their number of dimensions in which they are truly “nano”, their mode of hazard action, their solubility, fibrosity, or biopersistance^51^. Arts and coworkers^105^ presented a hazard based classification system for NPs with the aim to reduce testing necessity. According to this system, which resembles the cytotoxicity action terms of Brunner et al. ^51^, NPs are split up in four groups depending on their endpoint derived effects including intrinsic and system dependent variables. The critical properties are solubility (whereas solubility is defined as being dissolved before a maximum of 40 days in a biological system (*e.g.* lung)), fiber shape, which takes the special hazard deriving from respirable fibers into account, and general physico-chemical and biologic activity. Currently, risk assessment of engineered NPs in food deals currently mostly with Ag NPs for antibacterial purpose, TiO_2_ and silica as diffusion barriers, and nano-encapsulated ingredients^106^. Due to accumulation in the food chain nano-pesticides and veterinary medicines should be considered. Some concern, but yet no evidence, is that nanoencapsulated ingredients (*e.g*. micelles) are transported directly into the bloodstream. Due to their size it is possible but the discussion point is about the particles being broken down and metabolized long before reaching the blood stream.

**Concerning regulatory affairs**, any new substance, nano or not and independent of further use, which is commercially (!) produced or sold in Europe must be approved according to REACH (Registration, Evaluation, Authorisation and Restriction of Chemicals, (EG) Nr. 1907/2006 (REACH)). For approval a technical dossier concerning safe handling, exposure data, chemical, physical, toxicological, and ecotoxicological data (e.g. material safety data sheet) must be provided. If it comes to food, authorities currently still often consider NPs as new forms of old additives. The current status of the European Food Safety Authority (EFSA) is the general recommendation of *in vitro* and *in vivo* studies of NPs which are used in food^107^. There is also a recommendation not to use Ag NPs to extend shelf-life, and parallel a limitation to 50 μg Ag^+^/kg food, a concentration which has no biocide effect any more. FDA (Food and Drug Administration, USA) approves use of Ag in bottled water to a maximum of 17 μg Ag+/kg^108^, while use of TiO_2_ (nano or not) is currently without any restrictions^109^. Until 2013 there were 33 NP containing drugs approved and meanwhile FDA included NP size disclosure as an optional part of the approval process. In most cases the NP is part of the drug delivery system, not the drug itself^110^: *e.g.* antibody-drug conjugates, a nano-liposome based delivery platform, and albumin NPs. Other NPs are approved as imaging agents for diagnosis or additives to implant materials (*e.g.* bone cement).

### Workflow of NP hazard assessment

On cellular level the most important hazards concerning NPs are the release in form of toxic ions and molecules, oxidative stress with its' side effects like DNA damage, and cytoskeleton impairment. On the organism level the hazard will depend on the way and amount of uptake but the most likely occurring effects would be inflammation, immune system activation or depression, and blood coagulation impairment. To assess them systematically, without wasting time, money and animal life and without overlooking less abundant but important effects we propose a workflow starting with predictive toxicology and modeling of NP properties followed by experimental confirmation. -Omics approaches are here highly recommended, not only to save time, cost and animal life but to check for other non-typically activated pathways, too. By random checks of the status of a few thousand transcripts or metabolites it is less likely to overlook an important pathologic mechanism activated by NPs.

A workflow for NP hazard assessment may therefore look like this:

1. Preparation by data analysis and modeling attempts: Comparison of the new material to existing NP data to get an idea what can be expected from this new NP especially in terms of uptake, ADME, activated biological pathways and expected pathology mechanisms.
2. A primary classification could include information about expected uptake rates, biopersistance and release of ions/molecules, fiber shape, biologic activity, surface activity, respectively, and a preliminary advice of safe handling. This information should lead the way to experimental confirmation.
3. Systematic design of experiment (DoE) to confirm the predictive model. The experiments should be designed to proof and quantify the expected uptake rates, the expected biological effects and to give an unbiased overall picture of the NPs impact to avoid overlooking unexpected mechanisms.

a. Experimental read-out should cover the most abundant NP effects *in vitro* (ion/molecule release, oxidative stress, DNA damage and cytoskeleton impairment) and *in vivo* (inflammation, immune system activation/depression, and blood coagulation) and give information about impact, quantity and kinetics of the observed effects.
b. To avoid overlooking of unexpected effects omics approaches should be used.
c. Different dose, exposure and uptake scenarios:

i. low dose and high dose exposure
ii. single and repeated dose, and chronic exposure studies
iii. respiratory and gastro-intestinal uptake (nasal, ocular, dermal, too, if appropriate)
4. The final classification should be performed according to the expected and proven data from 1 and 3.

**Figure 2:**
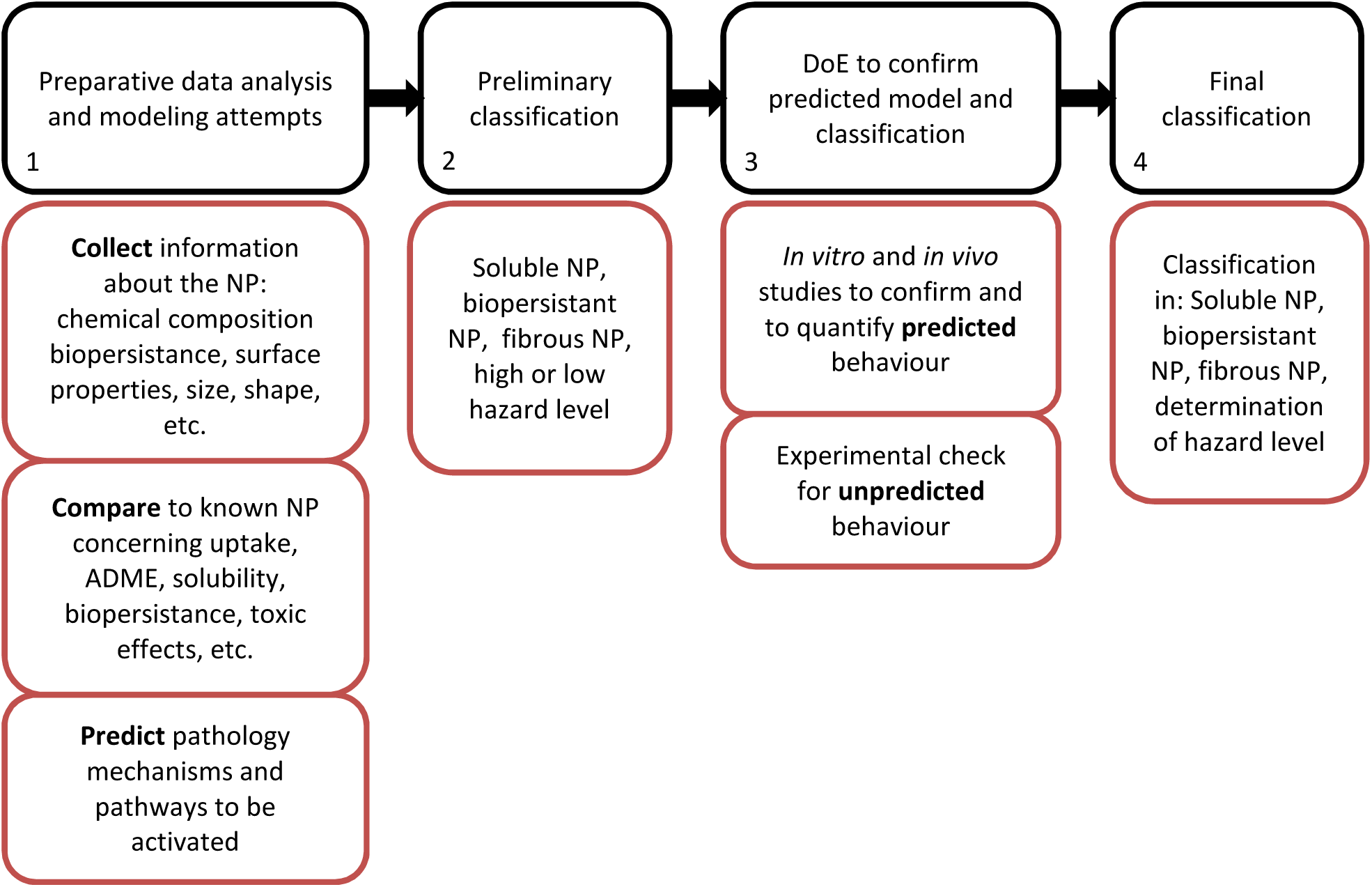
A workflow proposed for NP hazard assessment for new NPs: Collecting and comparing data about known NPs leads to prediction of potential hazards and preliminary classification. Carefully designed experiments shall test whether predicted and measured behavior match and support the final determination of hazard level.

## 5. Conclusion

In this review, we summarized the current status of knowledge concerning more or less abundant NP effects *in vitro:* and *in vivo,* namely ion toxicity, oxidative stress, inflammation, and fiber effects. We showed that omics approaches contribute to a better understanding of biological pathways activated by NPs, revealing not only the effects expected but giving an unbiased overview of changed metabolism, adverse pathway activation and gene signaling. We highlighted the possibilities to predict toxicological effects using modeling approaches and proposed an advanced workflow to assess NP hazard to improve risk assessment.

